# Towards a “Treadmill Test” for Cognition: Reliable Prediction of Intelligence From Whole-Brain Task Activation Patterns

**DOI:** 10.1101/412056

**Authors:** Chandra Sripada, Mike Angstadt, Saige Rutherford

## Abstract

Identifying brain-based markers of general cognitive ability, i.e., “intelligence”, has been a longstanding goal of cognitive and clinical neuroscience. Previous studies focused on relatively static, enduring features such as gray matter volume and white matter structure. In this report, we investigate prediction of intelligence based on task activation patterns during the *N*-back working memory task as well as six other tasks in the Human Connectome Project dataset, encompassing 19 task contrasts. We find that whole brain task activation patterns are a highly effective basis for prediction of intelligence, achieving a 0.68 correlation with intelligence scores in an independent sample, which exceeds results reported from other modalities. Additionally, we show that tasks that tap executive processing and that are more cognitively demanding are particularly effective for intelligence prediction. These results suggest a picture analogous to treadmill testing for cardiac function: Placing the brain in an activated task state improves brain-based prediction of intelligence.

## 1 Introduction

In addition to particular abilities associated with individual cognitive tasks, there is substantial evidence for an overarching general ability involved in performance across a diverse range of tasks.^1–5^ Intelligence tests composed of multiple subtests can yield accurate estimates of this general ability, usually denoted *g*, and which we here refer to as “intelligence”.^6, 7^ Intelligence is a fundamental dimension of individual differences and is a key contributor to a number of important academic, occupational, health, and well-being-related outcomes.^8–13^ There is thus substantial interest in understanding the neural basis of intelligence and in developing “neuromarkers” of intelligence, objective brain-based means of measuring individual differences.

One potentially promising route is to construct neuromarkers of intelligence from neuroimaging maps, including structural and functional imaging. Existing studies have mainly investigated brain size^14^, cortical thickness/gray matter volume^15, 16^, white matter structure^17^, and resting state functional connectivity^18–20^ (for reviews, see ^21–24^). A notable feature of these studies is they mainly examine stable, enduring features of the brain, features that are largely independent of the person’s current cognitive state, and in particular their actual exercise of the cognitive abilities that are relevant to intelligent task performance. An alternative approach for constructing neuromarkers of intelligence, whose rationale resembles that for cardiac treadmill testing, attempts to first place the brain in an activated state that engages these cognitive abilities, rendering brain features associated with these abilities more “visible” to functional imaging (see ^25^ for a suggestion along these lines).

Interestingly, while task-based neuroimaging studies have been extensively used to investigate the brain-basis of intelligence, they have thus far been used nearly exclusively for the purposes of localization: Activation patterns in higher versus lower intelligence individuals are compared to identify the places in the brain where there are statistically significant differences.^26–30^ Task-based studies have thus far not been used widely in a prediction framework in which whole-brain activation patterns are harnessed for making predictions of each subject’s intelligence.

In the current study, we address this notable gap. Utilizing the Human Connectome Project’s 1200 release, we construct a highly reliable measure of intelligence from 10 measures from the NIH Toolbox and Penn Neurocognitive Battery. We then examine brain-based prediction of intelligence from contrast maps derived from the *2*-back working memory task as well as six other fMRI tasks (19 task contrasts in total), and we establish two things. First, task-based activation patterns allow highly reliable prediction of intelligence, with performance appreciably higher than that reported in other neuroimaging modalities. Second, tasks that tap executive processing and that are more cognitively demanding yield more accurate predictions of intelligence.

## 2 Methods

### 2.1 Subjects and Data Acquisition

All subjects and data were from the HCP-1200 release^31, 32^ and all research was performed in accordance with relevant guidelines and regulations. Subjects provided informed consent, and recruitment procedures and informed consent forms, including consent to share de-identified data, were approved by the Washington University institutional review board. Subjects completed two runs each of seven scanner tasks across two fMRI sessions, using a 32-channel head coil on a 3T Siemens Skyra scanner (TR = 720ms, TE = 33.1ms, 72 slices, 2mm isotropic voxels, multiband acceleration factor = 8) with right-to-left and left-to-right phase encoding directions. Comprehensive details are available elsewhere on HCP’s overall neuroimaging approach^31, 33^ and HCP’s task fMRI dataset^34^.

Subjects were eligible to be included if they had available MSMAll registered task data for both runs of all seven tasks, had full behavioral data, and no more than 25% of their volumes in each run exceeded a framewise displacement threshold of 0.5mm. These exclusions resulted in 958 subjects. This was further reduced to 944 subjects in generating the Train/Test set split (see below).

### 2.2 Data Preparation

Data was preprocessed through the HCP minimally preprocessed pipeline, which is presented in detail by Glasser *et al.*^35^ Briefly, the pipeline includes gradient unwarping, motion correction, fieldmap distortion correction, brain-boundary based linear registration of functional to structural images, non-linear registration to MNI152 space, and grand-mean intensity normalization. Data then entered a surfaced-based preprocessing stream, followed by grayordinate-based processing, which involves data from the cortical ribbon being projected to surface space and combined with subcortical volumetric data.

### 2.3 FMRI Tasks

We used contrasts from seven HCP tasks, described in brief in Table 1 (detailed descriptions are available elsewhere^32, 34^).

**Table 1:** Seven HCP fMRI Tasks.

|  |  |
| --- | --- |
| *N-back task* | Participants respond when the picture shown on the screen is the same as the one two trials back (=2-back condition) or the same as one shown at the start of the block (=0-back condition). |
| *Incentive Processing* | Participants guess whether the number on a mystery card will be more or less than 5 and win or lose money (*reward* condition = mostly wins; *loss* condition = mostly losses) |
| *Motor* | Participants move fingers, toes, and tongue |
| *Language Task* | Participants answer questions about Aesop's fables (=story condition) or math problems (=math condition). |
| *Social Cognition Task* | Participants watch video clips of objects interacting in an agentive way (=theory of mind condition) or random way (=random condition). |
| *Relational Task* | Participants identify the dimension along which a cue pair of objects differs and determine if a target pair differs along same dimension (=relational condition). Or they determine if a cue object matches a member of a target pair along a given dimension (=match condition). |
| *Emotion Task* | Participants decide whether one of two presented faces match one at the top of the screen (=face condition) or else they perform the same task with shapes (=shape condition) |

At the single subject-level, fixed-effects analyses were conducted using FSL’s FEAT to estimate the average effects across runs within-participants, using 2mm surface smoothed data. Some tasks permitted multiple contrasts beyond the standard experimental versus control condition (e.g., *N*-back allows additional contrasts based on all four stimulus types). To reduce the complexity of the analysis and avoid loss of power from smaller number of trials, we focused on the standard contrasts yielding a total of 19. A full list of filenames of the contrast maps used can be found in the Supplemental Table S1.

### 2.4 Constructing a General Intelligence Factor

We conducted an exploratory factor analysis utilizing the strategy and associated code supplied by Dubois and colleagues (https://github.com/adolphslab/HCP_MRI-behavior), who recently investigated prediction of intelligence from resting state fMRI in the HCP dataset^36^. Unadjusted scores from ten cognitive tasks for 1181 HCP subjects were included in the analysis (subjects with missing data or MMSE < 26 were excluded), including seven tasks from the NIH Toolbox (Dimensional Change Cart Sort, Flanker Task, List Sort Test, Picture Sequence Test, Picture Vocabulary Test, Pattern Completion Test, Oral Reading Recognition Test) and three tasks from the Penn Neurocognitive Battery (Penn Progressive Matrices, Penn Word Memory Test, Variable Short Penn Line Orientation Test), with additional details supplied in^36^.

We applied Dubois and colleagues’ code to this data, which in turn uses the omega function in the psych (v 1.8.4) package^37^ in R (v3.4.4). In particular, the code performs maximum likelihood-estimated exploratory factor analysis (specifying a bifactor model), oblimin factor rotation, followed by a Schmid-Leiman transformation^38^ to find general factor loadings.

To assess reliability, in a separate analysis, we re-ran the factor analysis excluding 46 subjects that had Test/Retest sessions available. We then estimated factor scores for both sessions for these subjects and calculated test/retest reliability via intraclass correlation (we used ICC(2,1) in the Shrout and Fleiss scheme^39^).

### 2.5 Train/Test Split

The 958 subjects after exclusions were divided into two groups. To generate a test dataset of 100 unrelated subjects we first selected the 86 subjects with no siblings who passed exclusion criteria. Then, we randomly selected 14 families from those families with 2 siblings included and randomly dropped one sibling of each pair. This resulted in 944 total subjects for our analyses, with 844 in the training data and 100 unrelated subjects in the test set.

### 2.6 Dimensionality Estimation and Principal Component Analysis

In the training dataset, each subject’s unthresholded task data from 19 task contrasts was vectorized and concatenated yielding 19 matrices, each with dimensions of 844 subjects by 91282 grayordinates. We next estimated the number of intrinsic dimensions associated with each task contrast. This was accomplished by submitting each of the 19 subjects x grayordinates contrast matrices to the dimensionality estimation procedure of Levina and Bickel^40^. This is a maximum likelihood estimation method based on distance between close neighbors, which we previously successfully applied to HCP resting state data^41^.

Dimensionality estimation found a mean of 72 dimensions across the 19 task contrasts. Because prior studies by our group^41^ showed small differences in the number of components make little difference in classifier performance, we chose the round number of 75 components for each task. We also verified that using the exact task-specific number of components made no difference to the outcome (see Supplemental Table S2).

Next, each of the 19 subjects x grayordinates contrast matrices was submitted to principal components analysis using the pca function in MATLAB (2015b), yielding, for each contrast, 843 components ordered by descending eigenvalues.

### 2.7 Brain Basis Set Modeling

Our aim was to predict each subject’s intelligence from their expression scores for *n* components (where *n* was typically set to be 75; see above). To accomplish this, we used Brain Basis Set (BBS) modeling, previously described in detail^41^.

In a training dataset, we calculate the expression scores for each of the *n* components for each subject by projecting their data onto the 75 principal components. We then fit a linear regression model with these expression scores as predictors and the phenotype of interest (i.e., intelligence) as the outcome, saving **B**, the *n x 1* vector of fitted coefficients, for later use. In a test dataset, we again calculate the expression scores for each of the *n* components for each subject. Our predicted phenotype for each test subject is the dot product of **B** learned from the training dataset with the vector of component expression scores for that subject. We assessed performance of BBS-based prediction of intelligence by calculating the correlation between predicted versus actual intelligence in the test sample.

### 2.8 Addressing Additional Potential Confounds

In an additional analysis, we used multiple regression to remove a number of potential confounds from the intelligence variable (i.e., *g*). Similar to Dubois et al.^36^, variables regressed were: age, handedness, gender, brain size, multiband reconstruction algorithm version number (HCP variables: Age_In_Yrs, Handedness, Gender, FS_BrainSeg_Vol, fMRI_3T_ReconVrs), and mean framewise displacement (task-specific values were used). Analyses involving intelligence prediction were then repeated with the confound-cleansed variable. Results were broadly similar to the original analyses and are presented in Supplemental Table S2.

### 2.9 Consensus Component Maps for Visualization

We used BBS with 75 whole-brain components to make predictions about intelligence. To help convey overall patterns across the entire BBS predictive model, we constructed “consensus” component maps. We first multiplied each component map with its associated beta from the fitted BBS model. Next, we summed across all 75 components yielding a single map, and *z* scored the entries.

## 3 Results

### 3.1. Constructing a *g* Factor From Ten HCP Behavioral Tasks

As reported by Dubios and colleagues^36^, a bifactor model with a general factor *g* and four group factors, fit the data very well (CFI=0.990; RMSEA=0.0311; SRMR=0.0201; BIC=-0.519). The solution is depicted in Figure 1. Following Dubois and colleagues, we interpret the four group factors as: 1) Crystallized Ability; 2) Processing Speed; 3) Visuospatial Ability; and 4) Memory.

**Figure 1:**
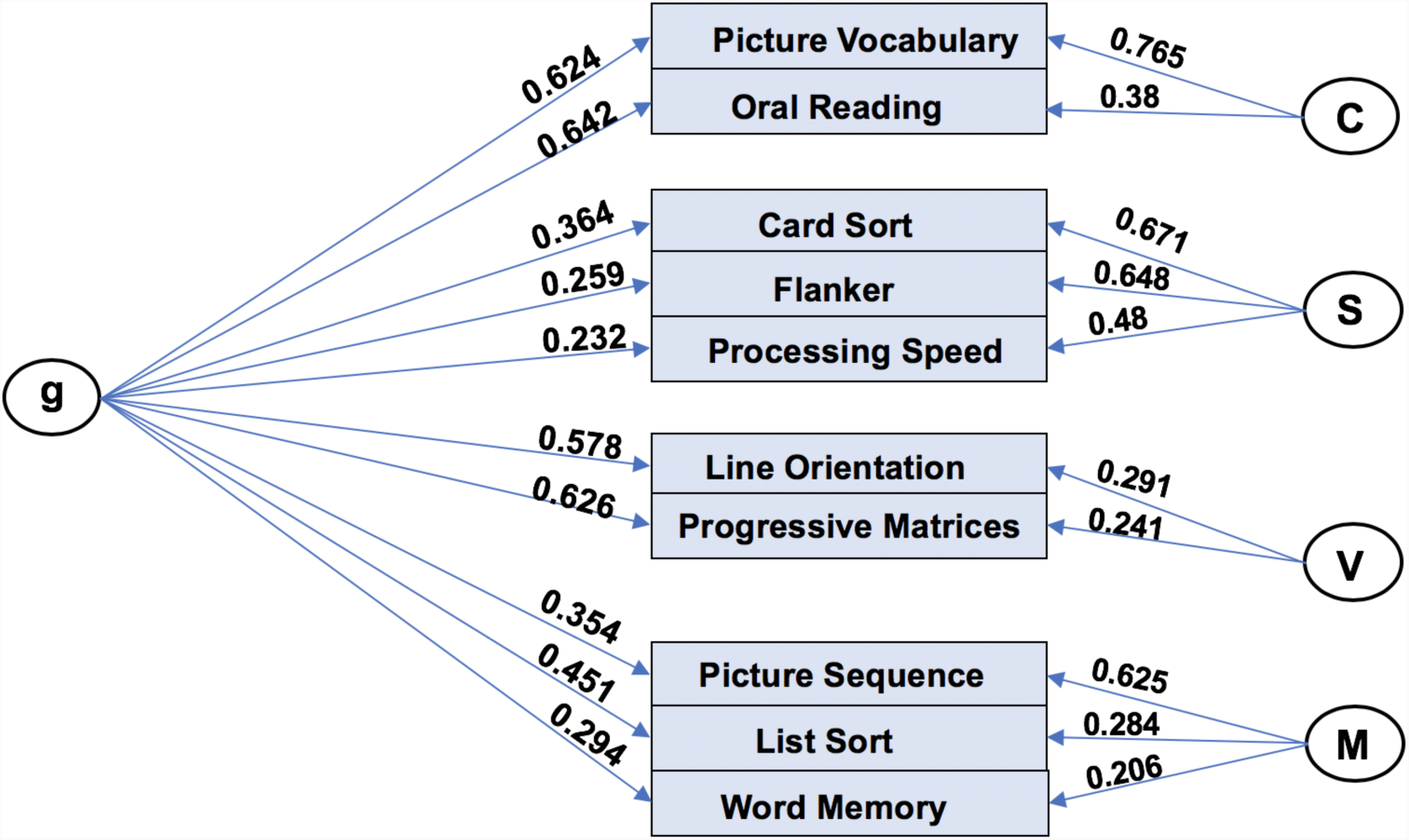
Bifactor Model Based on Ten Behavioral Tasks from the HCP Dataset with General Factor (“g”) and Four Group Factors. C=Crystallized Intelligence, S=Processing Speed, V=Visuospatial Ability, M=Memory.

The general factor *g*, which we refer to throughout as “intelligence” and which is the focus of this report, accounts for 58.5% of the variance (coefficient omega hierarchical ω^42^), while group factors account for 18.2% of the variance. Based on the 46 subjects in the retest dataset for HCP, test-retest reliability for *g* was found to be, 0.78, which is conventionally classified as very good (we used ICC(2,1) in the Shrout and Fleiss scheme^39^).

### 3.2 Contrasts associated with the *N*-Back task are highly effective at predicting intelligence in out-of-sample subjects

Because working memory has been strongly and consistently linked with intelligence^35, 43, 44^, we first investigated prediction of intelligence based on the *N*-back working memory task. A BBS model utilizing 75 components was fit in the training dataset and then applied to the test dataset. The correlation in the test dataset between predicted intelligence and actual intelligence was 0.68 (*p* = 8.5 × 10^−15^).

Figure 2 shows the top three components based on statistical significance displayed so that greater expression of these components predicts greater intelligence. These components include large activations in supplementary motor area (SMA), precuneus, and dlPFC, as well as deactivations in anterior default mode network (DMN). To convey “average” patterns across all 75 components, we constructed consensus component maps (see Methods) and they are displayed in Figure 3. These show additional patterns predictive of intelligence, including deactivation of posterior cingulate cortex and fronto-polar cortex.

**Figure 2:**
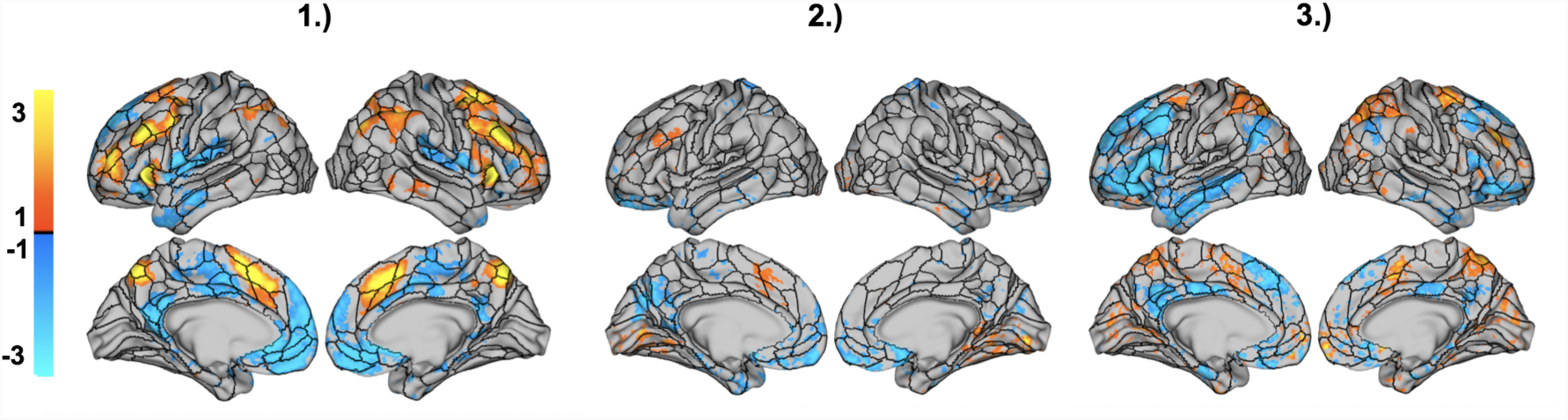
Visualization of the Three Most Intelligence-Predictive Components From the 2-back vs. 0-back Task Contrast. The three most statistically significant components are shown from a 75-component brain basis set model trained to predict intelligence scores.

**Figure 3:**
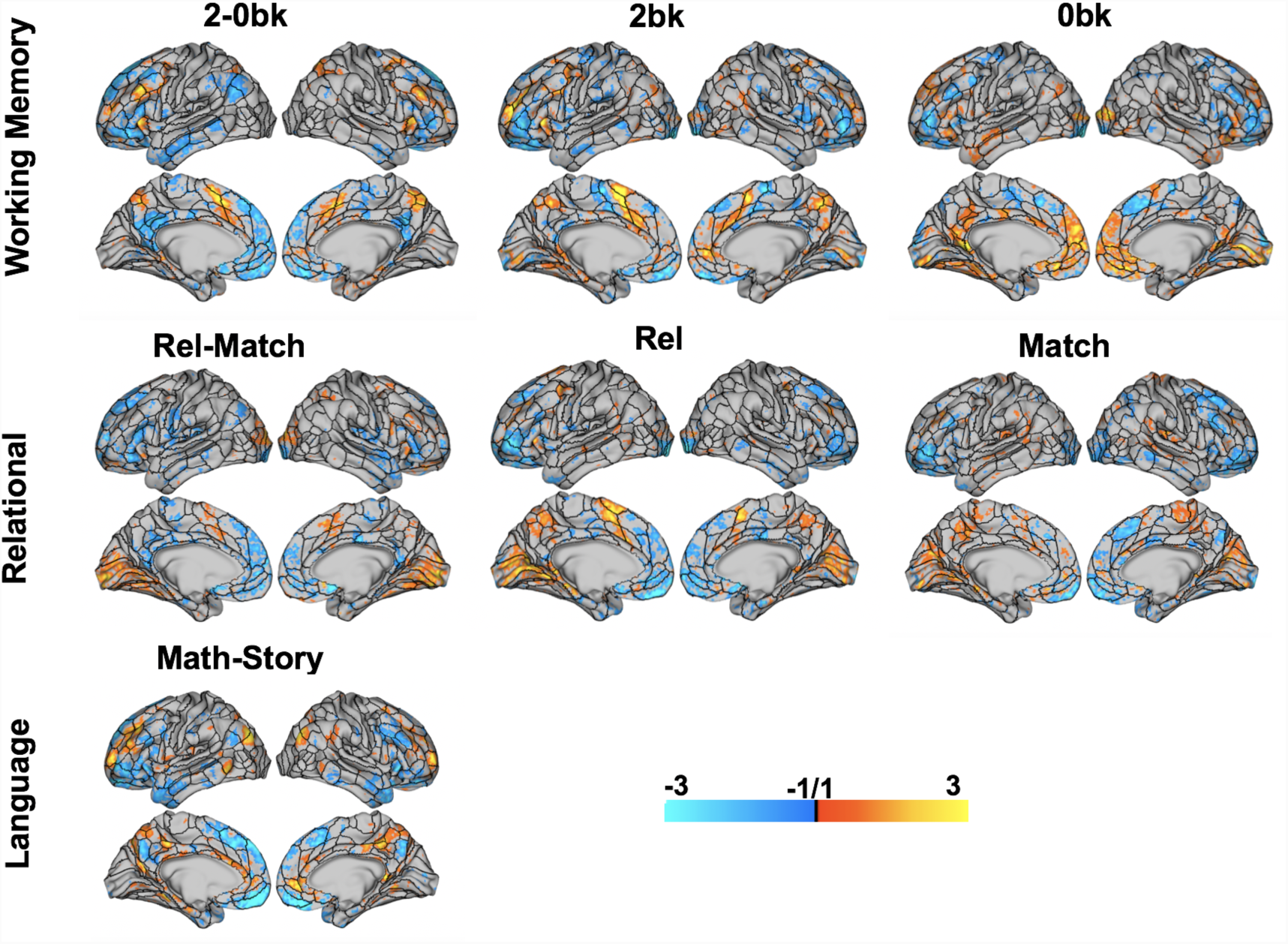
Consensus Component Maps for Seven Task Contrasts Highly Predictive of Intelligence. Each consensus component map captures aggregate patterns across a 75-component brain basis set model for prediction of intelligence. *(top row) N-Back Task; (middle row) Relational Task; (bottom row) Math – Story Contrast.*

We next trained additional BBS models on the *2*-back vs. baseline and *0*-back vs. baseline contrasts. The correlation in the test dataset between predicted intelligence and actual intelligence was 0.58 (*p* = 2.0×10^−10^) and 0.43 (*p* = 6.2×10^−6^), respectively. The consensus component models in Figure 3 revealed an interesting change in directionality across these contrasts. For example, pre-SMA strongly predicts greater intelligence in the *2*-back contrast vs. baseline but the reverse is true in the *0*-back vs. baseline contrast. Additionally, less activation (i.e., deactivation) of the anterior DMN predicts higher intelligence in the *2*-back vs. *0*-back contrast, but the reverse is true in the *0*-back vs. baseline contrast.

### 3.3 Looking across all 19 task contrasts, tasks involving executive processing and higher cognitive demand are more effective in predicting intelligence

We next compared intelligence prediction based on the three *N*-back task contrasts with the remaining 16 contrasts from the other six HCP tasks. As with the *N*-back task, we trained BBS models for each task contrast in the train dataset and applied the models to the held out test dataset.

Results are shown in Figure 3. Tasks involving executive processing were top performers, including contrasts from the *N*-back task and relational reasoning task, as well as the math vs. story contrast of the language processing task. The reward vs. baseline and punishment vs. baseline contrasts of the gambling task, both of which involve making numerical judgments, also performed well.

### 3.4 FPN/DMN activation, a measure of task demandingness, predicts which tasks are effective for intelligence prediction

A number of studies have observed that tasks that are cognitively demanding produce activation in regions of frontoparietal network (FPN)^45–48^ and deactivation of regions of default mode network (DMN)^49–52^. Building on these observations, we hypothesized the more cognitively demanding tasks (operationalized in terms of activation levels of FPN and DMN) should be more effective in predicting intelligence. To ensure comparability across task contrasts, we focused on the 12 task contrasts that compared a task condition versus resting baseline. We performed a regression analysis with accuracy of intelligence prediction as the outcome variable and FPN and DMN activation as predictors. Results, shown in Figure 4, support our hypothesis: The correlation across task contrasts between predicted accuracy, based on FPN and DMN activation, and actual accuracy in predicting intelligence is 0.67 (FPN standardized β = 0.63, DMN standardized β = - 0.58, *p* = 0.0482).

**Figure 4:**
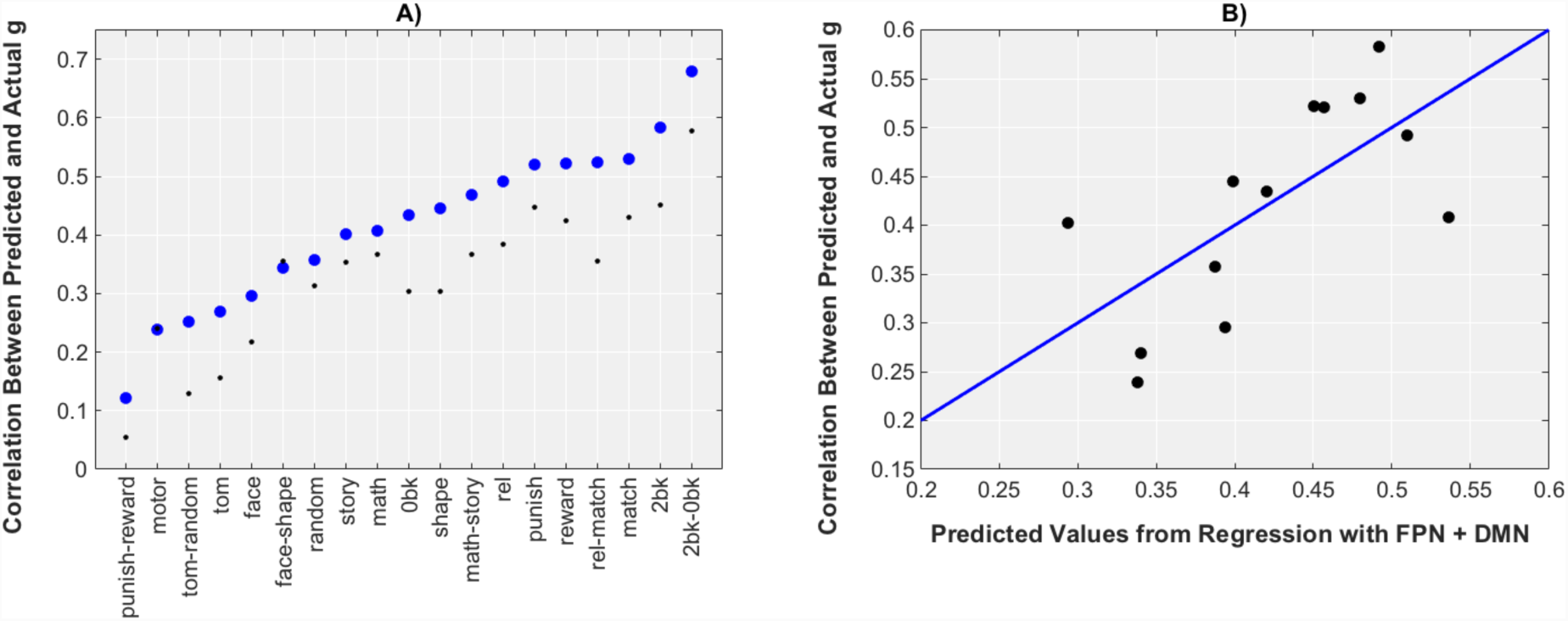
Intelligence Prediction Across 19 Task Contrasts. A. Prediction of Intelligence Using Brain Basis Set Modeling For 19 HCP Task Contrasts. Small Black Dots = results with additional regression of a number of potential confounds (see §2.8). B. FPN/DMN Activation Patterns During a Task Strongly Predict Effectiveness of That Task in Predicting Intelligence.

### 3.5. Across the 19 task contrasts, activation signatures of intelligence are spatially distributed and task-specific

We next compared the consensus component maps associated with the 19 contrasts (seven maps are shown in Figure 3, and the remaining maps are shown in Supplemental Figure S1). Signatures of intelligence prediction associated with each task were highly distributed, with remarkable variation in these signatures across tasks. Prominently represented regions include: superior parietal cortex (reward vs. baseline, punishment vs. baseline), dlPFC (math vs. story), anterior insula (relational v. match), fronto-polar cortex (math vs. story), pre-SMA (relational vs. match), and visual cortex (relational vs. match, reward vs. baseline, punishment vs. baseline). All 75 components as well as consensus component maps for each of the 19 task contrasts have been shared on BALSA, the Human Connectome Projects’ website for sharing and hosting neuroimaging datasets and can be accessed here: https://balsa.wustl.edu/study/show/MZPv.

## 4 Discussion

This study is the first to systematically assess neuroimaging-based prediction of intelligence across multiple fMRI task conditions. We find that whole-brain task activation patterns are a highly effective basis for prediction of intelligence, with a model trained on activation during the *N*-back working memory task achieving a 0.68 correlation with intelligence scores in an independent sample. Additionally, we demonstrate that more cognitively demanding tasks are particularly effective for intelligence prediction. These results highlight the importance of placing the brain in an activated task state for accurate brain-based prediction of intelligence.

### Role of executive regions in prediction of intelligence

The importance of fronto-parietal network, as well as related executive regions (e.g., dorsal anterior cingulate), for intelligence has been highlighted in previous work, especially in Jung and Haier’s influential fronto-parietal integration theory^21^. In a similar vein, Duncan and colleagues proposed that “multiple demand” cortex— regions of the brain that activate across a broad range of cognitively demanding tasks^45^—are a primary substrate of intelligence^27^. The present study extends these findings by demonstrating a key role for fronto-parietal regions as a major source of discriminative information for making subject-level predictions of intelligence. This additional role of executive regions was supported in three complementary ways.

First, in looking across the set of 19 contrasts derived from seven HCP tasks, we found that tasks that tap executive processes were more predictive of intelligence (e.g., *N*-back task contrasts, relational reasoning task contrasts, and math vs. story contrast). Second, we found that FPN activation and DMN deactivation, highly associated with the cognitive demandingness of task conditions^45–52^, predict which task contrasts will be effective for intelligence prediction. Third, within highly predictive contrasts, such the *2*-back vs. *0*-back contrast and math vs. story contrast, activation patterns in executive regions were prominent among regions predictive of intelligence.

Interestingly, for certain regions, the directionality of prediction of intelligence exhibited some variability across task contrasts in a way suggestive of moderation by task difficulty (for example, see pre-SMA in *0*-back compared to *2*-back and in match compared to relational). These observations are consistent with a neural efficiency model of intelligence proposed by Neubauer and Fink^53^. They propose that higher intelligence is associated with greater processing efficiency in elementary cognitive tasks (leading to less activation in more intelligent individuals) but greater processing capacity in demanding cognitive tasks (leading to greater activation in more intelligent people), thus potentially explaining the flipped directions of activation observed across the easy and hard conditions of the *N*-back and other tasks.

While activation patterns in executive regions clearly play an important role in explaining the success of our task-based approach to intelligence prediction, there is still clear evidence for substantial discriminative information about intelligence located outside executive regions. This is apparent in looking at the consensus component images in Figure 3 as well Figure S1 in the Supplement. Activations are observed in distributed regions of cortex, including non-executive regions such as visual cortex, lateral temporal cortex, and temporal pole, among other regions.

### Comparison of task-based prediction with other modalities

Previous studies using neuroimaging for prediction of intelligence have primarily used structural measures, for example cortical thickness^15, 16^ or white matter structure^17^, for reviews see ^21–23^. In terms of functional fMRI, recent studies have examined resting state connectivity patterns.^18–20^ In an important study by Dubois and colleagues^36^ that examined the same HCP 1200 dataset used in the present study, they found resting state connectivity patterns predicted intelligence with a correlation of 0.44 in cross-validated testing (see also a related finding from our group^41^). While impressive, this is still substantially less than the peak correlation of 0.68 achieved in the present study in out-of-sample testing.

There are two interrelated reasons why task-based fMRI might potentially offer more reliable prediction of intelligence than other modalities. The first appeals to the “treadmill testing” idea already mentioned: actively^34^ engaging in cognitive tasks has the potential to unmask critical intelligence-relevant features of the brain that are otherwise invisible in other modalities such as structural brain imaging. A second potential advantage of task-based methods is specificity. Tasks are constructed by their designers to target specific psychological processes, often with control conditions that subtract away contributions from auxiliary processes of no interest. This will tend to make classification more accurate as the feature set is culled of a sizable number of uninformative features.

### Future Directions

Results from this study open the door to promising additional lines of research. It is notable that we used the set of imaging tasks that were included in the HCP dataset. These imaging tasks, in turn, were selected based on diverse considerations (see ^34^), but maximizing prediction of intelligence was not among them. Thus, it is plausible that one can do still better: It should be possible to intentionally design and optimize an imaging task battery—based on findings from the literature as well as trial-and-error experimentation—to yield even more accurate task-based prediction of intelligence, and future work should explore this possibility.

In sum, this study firmly establishes the effectiveness of task-based fMRI for prediction of intelligence and demonstrates that tasks that tap executive processing and that are more cognitively demanding are associated with better prediction accuracy.

